# Frogs uncouple neural activity from oxygen consumption after hibernation

**DOI:** 10.64898/2026.02.03.703589

**Authors:** Hafsa Yaseen, Joseph M. Santin

**Author notes:** **Ethics approval statement:** Approval for the use of animals was approved by the Animal Care and Use Committee at the University of Missouri (protocol #39264). **Data availability statement:** The data that support the findings of this study are available from the corresponding author upon reasonable request.

## Abstract

**Aim:** Aerobic metabolism supplies ∼90% of the ATP for neural activity. In frogs, activity has large aerobic needs typical of an average vertebrate, but surprisingly, can shift to using only glycolysis upon emergence from hibernation. We hypothesized that hibernation triggers a global reduction in the aerobic cost of neural function.

**Methods:** We simultaneously measured activity of the brainstem respiratory network via motor nerves and tissue oxygen partial pressure (pO_2_) *in vitro* from control and hibernated bullfrogs (4 weeks cold submergence; 4°C). To identify which functions differentially consume O_2_, we sequentially blocked activity and various cellular processes requiring activity-independent ion regulation and used the resulting tissue pO_2_ change (ΔpO_2_) as an index of O_2_ consumed. We further assessed how activity varies as a function of tissue pO_2_ and how O_2_ consumption varies across network activity levels.

**Results:** Despite similar network activity levels, we provide three lines of evidence that hibernation reduces its aerobic requirement. First, hibernators consume less oxygen for baseline activity. Second, network output remains stable from baseline to anoxia, while moderate hypoxia disrupts controls. Finally, accelerating activity does not enhance oxygen consumption as in controls, but aerobic metabolism ultimately increases during seizure-like activity.

**Conclusion:** Hibernating frogs reduce aerobic needs for sustaining physiological levels of neural activity, revealing how they overcome the challenge of restarting motor circuits on the background of hypoxia during emergence from hibernation. More broadly, vertebrate neural circuits seemingly constrained by aerobic metabolism can exhibit substantial plasticity in the aerobic requirements for function.

**Practitioner points:**

1. Hibernation in bullfrogs reduces the oxygen consumed by neural activity while maintaining normal network output, demonstrating that the aerobic requirements of brain function are not fixed in the vertebrate brain.
2. After hibernation, brainstem motor circuits maintain stable function from high levels of O_2_ to anoxia and do not increase O_2_ consumption when activity is elevated within the physiological range.
3. These findings reveal that vertebrate neural circuits can enter metabolic states requiring far less aerobic respiration, which may inform strategies for improving metabolic resilience in the brain during conditions of impaired oxygen delivery and other metabolic dysfunction.

## INTRODUCTION

Brain function is energetically expensive^1,2^. At the circuit level, synaptic transmission and the restoration of ion gradients following action potentials have high ATP demands, the majority of which is supplied by oxidative phosphorylation^3,4^. Given the ATP yield of aerobic respiration (30-32 moles ATP/1 mole glucose) compared to glycolysis (2 moles of ATP/1 mole of glucose), neural activity and oxygen consumption are tightly coupled, whereby increases in activity drive aerobic metabolism to sustain neuronal performance^5–7^. This relationship is especially critical, as disruptions to aerobic metabolism impair neural output and lead to neurological disease^8^. The requirement for aerobic respiration is conserved across most vertebrate nervous systems and has shaped prevailing views that sustained neural computation requires oxygen consumption^9^.

Despite the view that neural activity’s large energy demands remain fixed, the American bullfrog (*Aquarana catesbeiana*) provides a unique opportunity to address how neural circuits can enter states that operate with far less energy. Under normal circumstances frog circuits exhibit typical metabolic demands and rely heavily on oxidative metabolism^10,11^. However, activity of brainstem motor circuits that generate breathing improve function during anoxia from ∼10 minutes to upwards of 4 hours under otherwise identical experimental conditions after frogs emerge from hibernation^12,13^. This represents a striking display of metabolic plasticity, as this network has high metabolic demands, with rhythm generating circuits that use glutamatergic synaptic transmission, which is thought to incur large energy costs, and motor pools with neurons that fire action potentials at rates >100 Hz during rhythmic bursting^14–16^. This form of metabolic plasticity is physiologically significant because arterial pO_2_ can fall as low as 1–3 mmHg during submergence^17^. While low pO_2_ does not present a problem due to low metabolic rates and decreased need for many behaviors in the cold, the problem is much more apparent upon emergence. Indeed, since neural activity typically has oxygen demands, frogs emerging from hibernation must resume neural activity under conditions when pO_2_ is expected to be similarly low^18^, which would be difficult if they did not become able to function with a lower requirement for aerobic ATP production.

Mechanistically, there are two non-mutually exclusive possibilities that could explain how hibernation alters the brainstem to function with less O_2_: increasing ATP supply during hypoxia and/or reducing the aerobic metabolism required to sustain activity. First, while glycolysis produces ∼15-fold less ATP than oxidative phosphorylation, hypoxia can increase glycolysis through the Pasteur effect. Thus, hibernation may lead to cellular changes that bolster flux through glycolysis^19^ to sustain neural activity without oxygen. On the other hand, hibernation may reduce the dependence of neural activity on aerobic metabolism making while maintaining similar network output. While there are several ways neural circuits could in principle enhance their energy efficiency^20–22^, hibernation reduces the Ca^2+^ permeability of NMDA-glutamate receptors and enhances receptor desensitization^23^, which is likely to reduce the energetic burden of ion regulation at synapses. Resolving these possibilities may provide new insight into how neural circuits can shift between states with vastly different aerobic requirements while maintaining similar function, as well as leading to new ideas for improving metabolic resilience in the brain.

In this study, we assessed the extent to which hibernation alters activity’s aerobic requirements in the brainstem respiratory network. To estimate the aerobic metabolic cost of activity under normal (control) and hypoxia-tolerant (after hibernation) network states, we simultaneously measured network activity through motor nerve rootlets and tissue oxygen partial pressure (pO_2_) within the rhythm-generating circuit *in vitro*. This approach allows us to estimate aerobic metabolism because tissue pO_2_ reflects consumption by mitochondria without the confound of cerebral blood flow, where pO_2_ decreases relate to greater rates of aerobic metabolism and increases reflect less oxygen consumption^5,24^. We then assessed which cellular processes alter their energy requirements, the relationship between tissue pO_2_ and activity, and how oxygen consumption varies over a range of network intensities. Collectively, results here indicate that hibernation uncouples physiological activity from its typical aerobic requirement, demonstrating that the same neural circuit within a single species can enter states that require less aerobic respiration while upholding physiological function.

## RESULTS

The amphibian respiratory network *in vitro* generates output that occurs as rhythmic bursts, which are periodically interrupted by high-drive breathing events termed episodes^25^. While the electrochemical sensor used in this study could not resolve O_2_ changes associated with single bursts which last ∼1s, when breathing episodes spanned several seconds, pO_2_ transients of a few mmHg occurred in phase (Figure 1A). This supports the detection of oxygen consumption within or around the active respiratory network. Despite having similar baseline network outputs (Figure 1B), hibernators had higher average tissue pO_2_ across the range of depths from the surface of the brainstem to 350 µm at the baseline external O_2_ levels (Figure 1C), and also in 6 additional preparations that were measured down to 650 µm near the center of the tissue (Supplemental Figure 1). Moreover, lowering the oxygen in the bathing solution decreased tissue pO_2_ as expected, but values were higher in hibernators at most fixed external pO_2_ (Figure 1D). These observations indicate a reduction in brainstem O_2_ consumption in hibernators when oxygen is present.

**Figure 1.**
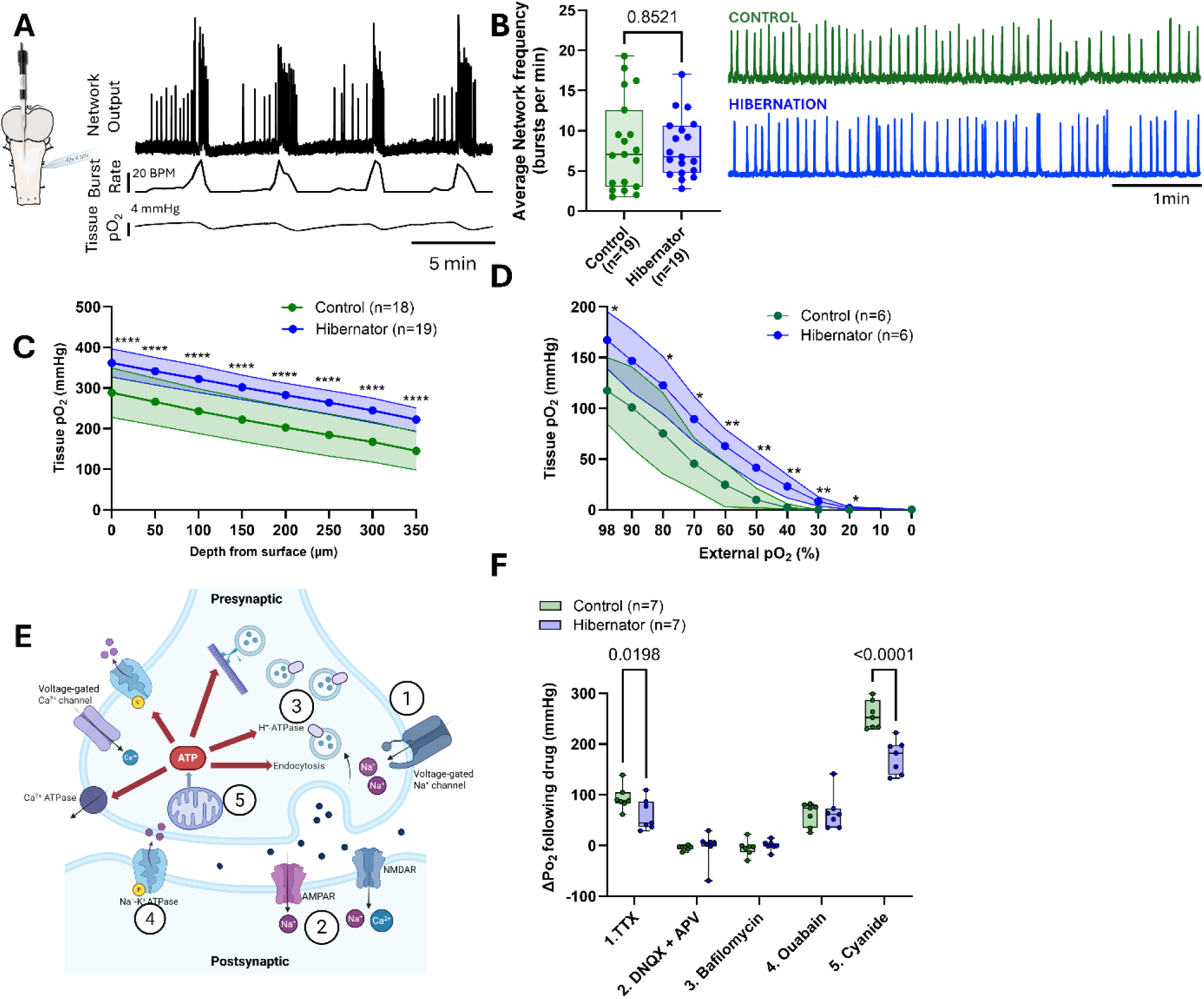
Hibernators have lower apparent oxygen consumption due to reduced energy demands of network activity and basal mitochondrial respiration. A. Schematic of the brainstem preparation with an oxygen sensor placed in the proposed rhythm-generating region. Example recording shows synchronized vagus nerve activity, network frequency, and corresponding tissue pO_2_ fluctuations. B. Control and hibernator brainstems showed similar baseline burst frequency (Welch’s unpaired t-test, p = 0.852; n = 19/group). C. Tissue pO_2_ depth profiles up to 350 µm showed significantly higher pO_2_ in hibernators (n=19) when compared to controls (n=18); two-way repeated measures ANOVA; main effect of group, F(1,35) = 31.18, p< 0.0001; main effect of depth, F(1.964, 68.74) = 590.9, p < 0.0001; depth x group interaction, F(7,245) = 0.4117, p = 0.8946; Holm–Šídák multiple comparisons test; ****p < 0.0001 at all depths. D. During gradual external pO_2_ reduction, hibernator brainstems maintained higher tissue pO_2_ throughout hypoxia induction (n = 6/group; two-way repeated measures ANOVA; main effect of group, F(1, 10) = 9.826, p = 0.0106 ; main effect of pO_2_, F(1,10)=145.2, p<0.0001; pO_2_ × group interaction, F(1.70, 11.7) = 4.929, p = 0.0426. Holm–Šídák multiple comparisons test, **p < 0.01, *p < 0.05). E. An illustration of major energy consuming pathways at the synapse with the blocked pathways numbered (made in Biorender). F. Summary of ΔpO_2_ after blocking voltage-gated Na^+^ channels, AMPA and NMDA receptors, vesicular ATPase, Na^+^-K^+^ ATPase and mitochondrial Complex IV (n = 7/group). Two-way repeated measures ANOVA revealed a significant main effect of group, (F(1, 12) = 122.3, p < 0.0001); main effect of drug, (F(4, 48) = 139.9, p< 0.0001 and drug × group interaction (F(4, 48) = 6.213, p = 0.0004). Hibernators showed significantly smaller ΔpO_2_ with TTX (p = 0.01) and sodium cyanide (p = 0.0003; Holm–Šídák multiple comparisons test).

We then used a pharmacological approach to identify which of the energy-consuming processes use less oxygen. Our rationale was to first block network activity and then apply a variety of antagonists that inhibit physiological processes involved in ion regulation that occur in the absence of ongoing network output. For this, we measured changes in averaged tissue pO_2_ during the sequential application of inhibitors (Supplemental Figure 2) that block network activity (tetrodotoxin; TTX), action potential-independent spontaneous excitatory postsynaptic potentials (i.e., “minis”; DNQX and APV), the vacuolar ATPase that pump H^+^ into vesicles (V-ATPase; bafilomycin), and the Na^+^-K^+^ ATPase that maintains resting ion gradients (ouabain), each of which has the potential to represent energy sinks in the brain as a result of activity or activity-independent housekeeping^22,26^ (Figure 1E). Finally, to account for remaining O_2_ consumption that would arise due to various other activity-independent physiological processes, we blocked complex IV of the mitochondrial respiratory chain (sodium cyanide).

In controls, TTX increased tissue pO_2_, demonstrating the consumption of O_2_ to power baseline activity. Blocking spontaneous excitatory transmission and then the V-ATPase had no further effects on tissue pO_2_. Inhibition of the Na^+^-K^+^ ATPase then raised tissue pO_2_, demonstrating a metabolic cost of regulating resting Na^+^ and K^+^ gradients. Poisoning mitochondria with cyanide led to a large increase in tissue pO_2_. When performing the same protocol on hibernators, less O_2_ was consumed by activity than controls, as TTX led to a significantly smaller rise in tissue pO_2_ (p = 0.0198). The other activity-independent processes consumed similar amounts of O_2_ as controls (DNQX+APV, p = 0.8966; bafilomycin, p = 0.5048; ouabain, p = 0.7841). However, hibernators showed a significantly smaller pO_2_ increase upon blocking mitochondrial Complex IV with sodium cyanide (p = 0.0003; Figure 1F). While these responses are presented as ΔpO_2_ before and after the sequential application of each drug, they are corroborated when analyzed absolutely (Supplemental Figure 2). Indeed, hibernators have a higher starting pO_2_ than controls, but no difference in TTX, indicating that TTX had a smaller effect in hibernators. After this, hibernators and controls had similar tissue pO_2_, indicating no differences across groups with respect to the O_2_ consumed by spontaneous synaptic postsynaptic events, the V-ATPase, and the Na^+^-K^+^ ATPase. Cyanide application had a smaller effect in hibernators, reaching lower absolute pO_2_ values. Finally bath O_2_ did not differ across groups. These results indicate that reduced oxygen consumption in hibernators is driven primarily by decreasing the cost of network activity and additional activity-independent mitochondrial O_2_ consumption that does not arise from the Na^+^-K^+^ ATPase, the V-ATPase, or spontaneous synaptic postsynaptic events.

Since hibernators had a reduced reliance on aerobic metabolism, we predicted that activity would be less sensitive to oxygen reductions and the associated decrease in aerobic metabolism. In control brainstems (n = 6), network activity decreased with tissue pO_2_ during graded reduction in bath O_2_. Each control preparation reached a critical tissue pO_2_ (p_crit_), beyond which the output either lost its rhythmicity or assumed a slow disrupted pattern of output while decreasing in average frequency (Figure 2A). In contrast, hibernators maintained normal activity throughout the entire range of graded hypoxia even at 0 mmHg (Figure 2B & 2C) with no apparent p_crit_ (Figure 2D). These results indicate that decreasing the reliance on aerobic metabolism allows hibernators to generate normal network activity across a wide range of tissue oxygen levels, from supraphysiological to anoxia.

**Figure 2.**
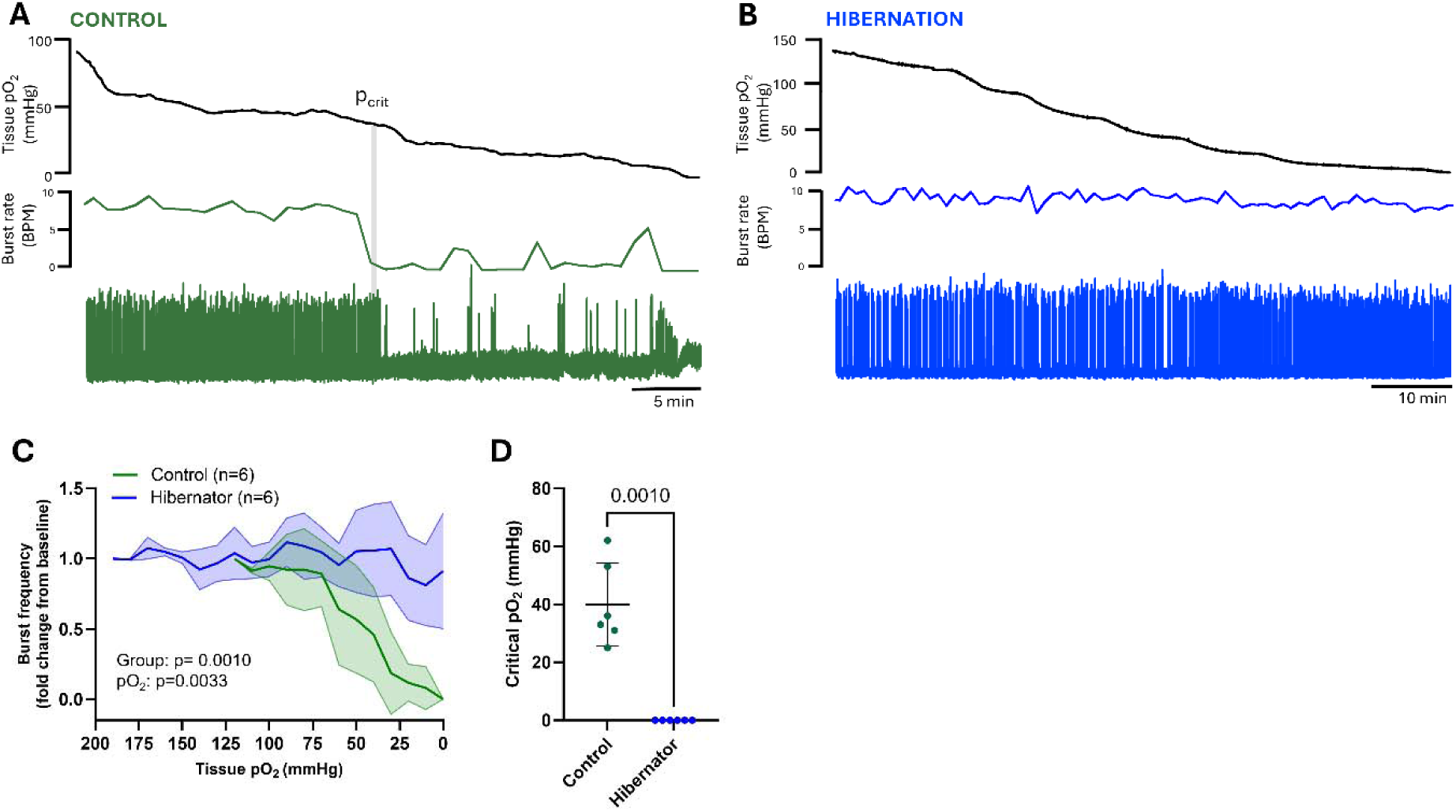
Brainstem motor activity no longer requires aerobic respiration after hibernation. A. Example of graded hypoxia experiment in a control, where nerve burst frequency declined strongly at a critical threshold (p_crit_). B. In hibernators (n = 6), neural activity remained stable even as tissue pO_2_ at ∼0 mmHg. C. Mean ± SD of fold change in burst frequency are shown for control and hibernator (n = 6 each) groups. Controls show a lower starting pO_2_ with an inflection point and subsequent decline in burst frequency as pO_2_ decreases, whereas hibernators have higher starting pO_2_ and remain relatively stable across the pO_2_ range. P values of group and pO_2_ from linear mixed model are shown in the figure. D. Critical pO_2_ thresholds: controls lost rhythmic activity at significantly higher pO_2_ than hibernators, which showed no apparent critical pO_2_ threshold as all preparations were active at 0 mmHg, and therefore assigned a p_crit_ value of 0 mmHg (Welch’s unpaired t-test, t(5) = 6.847, p = 0.001; n = 6 per group).

Beyond baseline activity, increased network activity drives oxygen consumption through synaptic vesicle cycling and restoration of ion gradients for action potentials and synaptic transmission, which is needed to sustain activity^3,7^. To test whether increased network activity would accelerate aerobic metabolism, we bath-applied the β-adrenergic receptor agonist, isoproterenol^27^. Example traces show that increases in activity were associated with a significant decline in average tissue pO_2_ in controls over the 30-minute application, indicating increased oxygen consumption to meet the demands of elevated activity (Figure 3A). In contrast, hibernators maintained a stable tissue pO_2_ despite achieving comparable increases in burst frequency (Figures 3B-3E). These results further support that metabolic adaptation in hibernators decouples neural activity from its typical aerobic support, allowing them to increase motor output without accelerating oxygen consumption.

**Figure 3.**
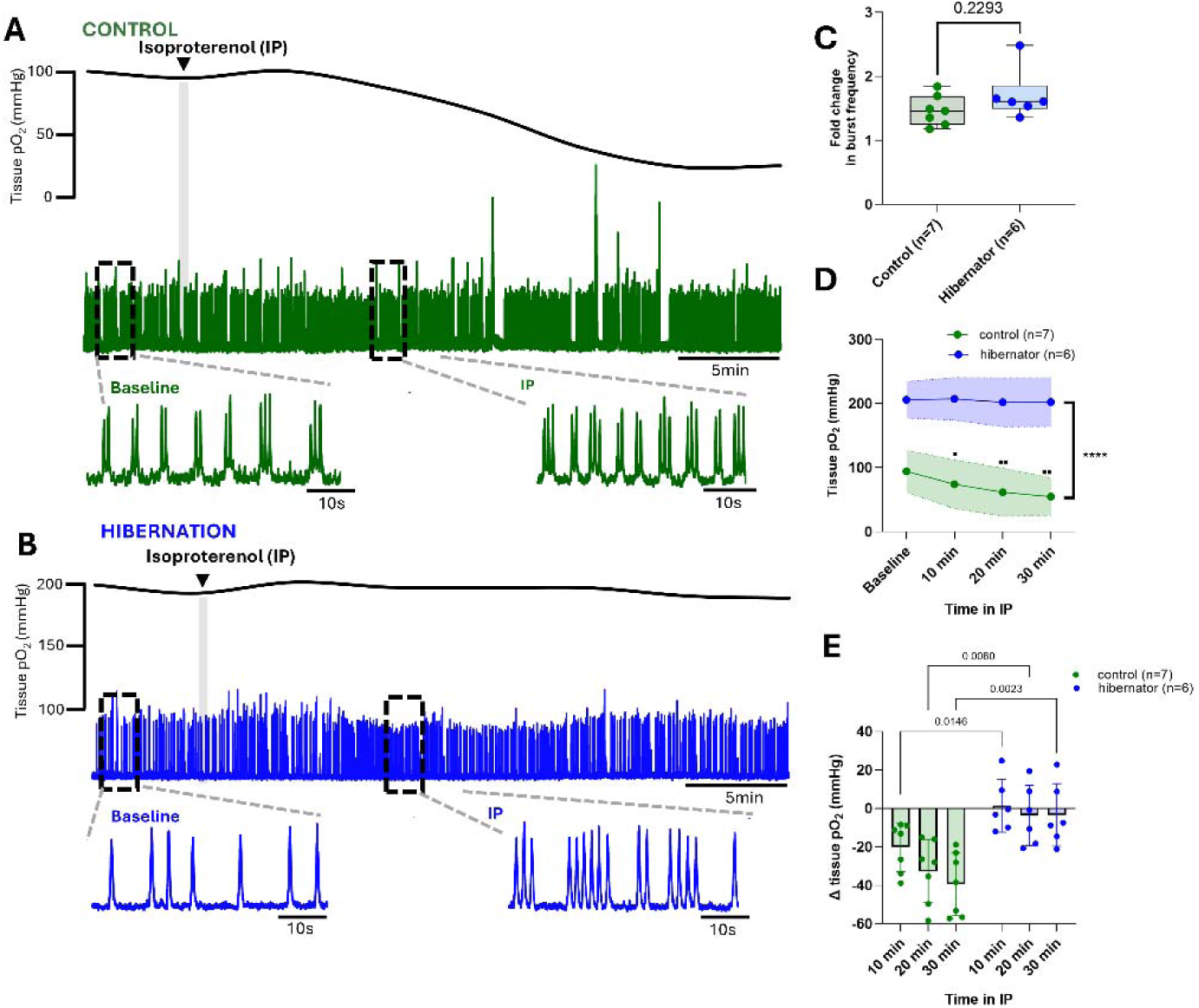
Hibernators sustain elevated neural activity without consuming more oxygen. A. Representative traces showing changes in tissue pO_2_ and vagus nerve activity during isoproterenol exposure in a control brainstem. Increased burst frequency caused a drop in tissue pO_2_. The insets show nerve activity of control brainstem at baseline and during isoproterenol exposure. B. Tissue pO_2_ in hibernators remained stable during elevated burst frequency. The insets show nerve activity of hibernator brainstem at baseline and during isoproterenol exposure. C. Fold change in burst frequency during isoproterenol application for controls (n=7) and hibernators (n=6; Welch’s unpaired t-test, t(7.931) = 1.302, p = 0.229). D. Absolute tissue pO_2_ at different timepoints of isoproterenol application (n = 7 controls, n = 6 hibernators). Two-way repeated measures ANOVA; main effect of group, F(1, 11) = 51.09, p < 0.001; main effect of time, F(2.038, 22.42) = 10.92, p = 0.0005; time × group interaction, F(2.038, 22.42) = 7.1, p = 0.0037. **** indicates p<0.0001 between controls and hibernators at all time points, and ▪▪p<0.01, and ▪ p<0.05 indicate differences from baseline within the control group (Holm–Šídák multiple comparisons test). E. ΔpO_2_ at 10, 20, and 30 minutes of isoproterenol application for controls and hibernators n = 6). Two-way repeated measures ANOVA; main effect of group, F(1, 11) = 16.2, p = 0.0020; main effect of time, F(1.338,14.71) = 4.9, p = 0.338; time × group interaction, F(2, 22) = 1.6, p = 0.2173; Holm–Šídák multiple comparisons test, **p < 0.01, *p < 0.05.

Finally, we asked if this metabolic decoupling extends to extreme hyperexcitability. We induced pathological network activity by blocking fast GABAergic inhibition with bicuculline, previously shown to generate high-amplitude bursts that resemble seizure-like activity in controls and hibernators^28^. In controls, bicuculline triggered abnormal bursts, which were accompanied by a large drop in tissue pO_2_, indicating substantial increases in aerobic respiration (Figures 4A, 4C-4D). Hibernators also exhibited an initial, large drop in pO_2_ following the first abnormal burst, demonstrating the ability to increase aerobic metabolism when activity goes beyond the physiological range (Figures 4B-4D). While the absolute decrease in pO_2_ from baseline was the same in controls (Δ92 ± 36.4 mmHg) and hibernators (Δ120 ± 22.4 mmHg; unpaired t-test, p = 0.31), after this initial consumption event, hibernators stabilized at a tissue pO_2_ approximately double that of controls (Figures 4B and 4C), indicating that aerobic metabolism was less under the same hyperexcitable conditions. Despite less oxygen consumption in the presence of bicuculine, hibernators maintained normal respiratory network output, while controls produced mainly seizure-like bursts (Figure 4A inset iii, 4D). Therefore, when pushed to the extreme, hibernators meet these demands by consuming O_2_ but ultimately maintain activity homeostasis during conditions of severe hyperexcitability with less aerobic metabolism.

**Figure 4.**
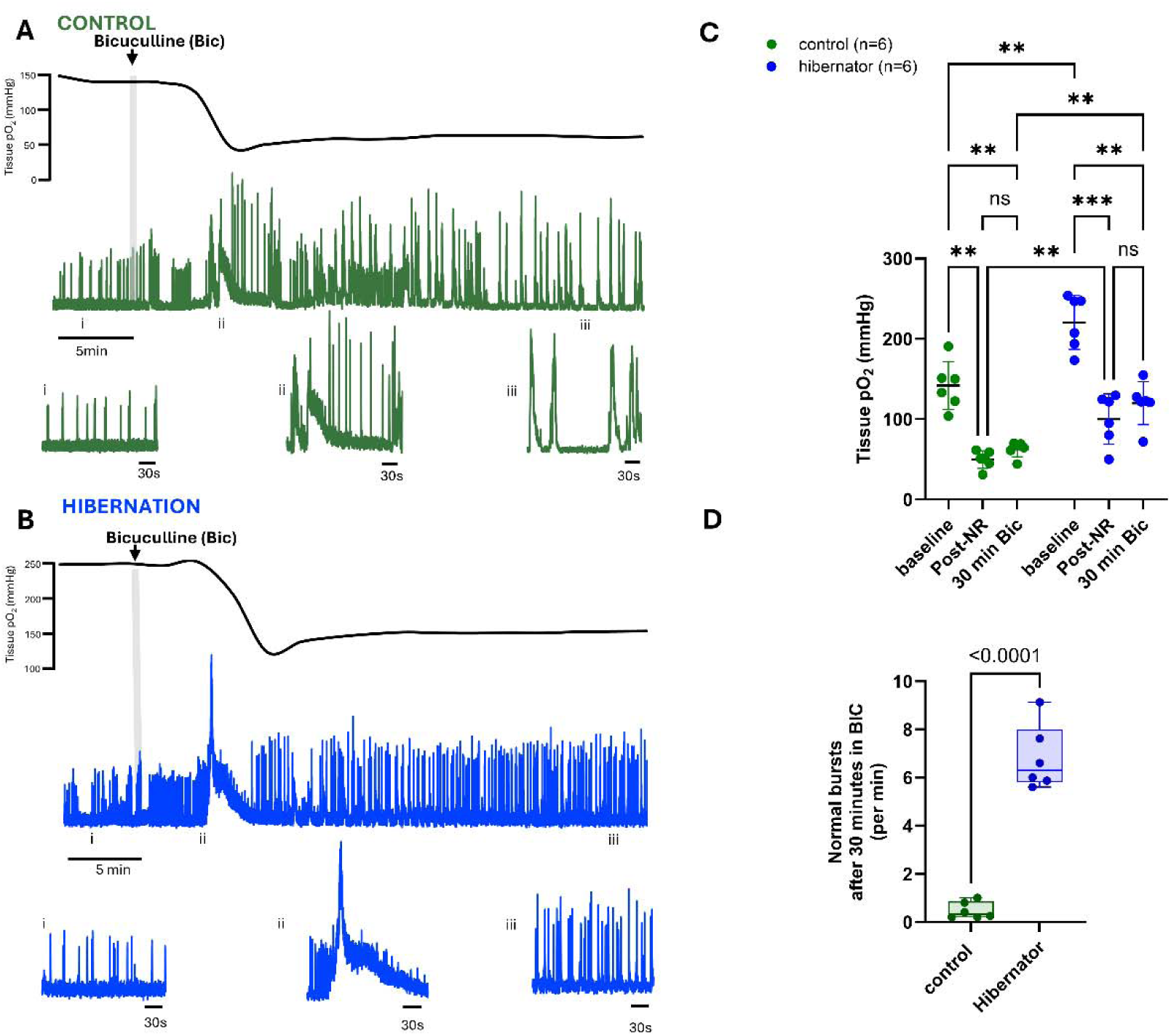
Hibernators rely on aerobic metabolism during pathological hyperexcitability but maintain elevated tissue oxygen. A. Representative recordings of tissue pO_2_ and vagus nerve activity in a control brainstem during application of the GABA_A_ receptor antagonist, bicuculline. Expanded nerve traces show baseline respiratory activity (RESP), the first large chaotic burst during bicuculline application, and then at the end of the 30 minute bicuculline exposure. B. Representative recordings of tissue pO_2_ and nerve activity in a hibernator brainstem during application of bicuculline. Expanded nerve traces show baseline respiratory activity (RESP), the first large chaotic burst during bicuculline application, and then at the end of the 30-minute bicuculline exposure. C. Absolute tissue pO_2_ at baseline, immediately after the first major non-respiratory burst (post-NR), and at the end of bicuculline treatment in control and hibernator brainstems (n = 6 per group). Both controls and hibernators face a significant drop in tissue pO_2_ post-NR, however tissue pO_2_ settles at a higher level in hibernators as compared to controls. Two-way repeated measures; main effect of group, F(1, 10) = 29, p = 0.0003; main effect of time, F(1.438, 14.38) = 39244, p < 0.0001; time × group interaction, F(2, 20) = 1.6, p = 0.2224; Holm–Šídák multiple comparisons test, ***p < 0.001, **p < 0.01. D. Number of normal bursts per minute after 30 minutes of bicuculline application in control and hibernator brainstem (n=6 per group; Welch’s unpaired t-test, t(5) = 11.15, p < 0.0001).

## DISCUSSION

Owing to the high ATP yield of aerobic respiration, increases in oxidative metabolism are thought to be essential for meeting the energy costs incurred by neural activity, from flies to mammals^6,29^. Energy limitations in neurological disorders have prompted interest in animal adaptations as sources of novel mechanisms for improving metabolic robustness in neural systems^18,30–33^. In frogs, hibernation transforms brainstem circuits to function during lack of oxygen by altering synapses such that glycolysis supports activity as a lone ATP source^12,13^. While impressive, we now show that neural activity can be uncoupled from its support by aerobic metabolism, even when oxygen is readily available. These results introduce that circuits with a seemingly obligate requirement for aerobic metabolism can enter a low-cost state that achieves the same functional output over a wide range of energetic challenges, including hypoxia, elevated activity, and pathological hyperexcitability, all while consuming less O_2_.

Based on the cellular processes addressed, neural activity, activity-independent Na^+^/K^+^ ATPase turnover, and additional housekeeping processes that are activity-independent are the main contributors of O_2_ consumption in the brainstem, with minimal input from spontaneous excitatory postsynaptic currents and the V-ATPase involved in filling vesicles with neurotransmitter (Figure 1F, Supplemental Figure 2). Despite similar levels of neural activity, hibernator brainstems exhibited a reduction in O_2_ consumption, driven by less O_2_ consumed by neural activity and less spent on other activity-independent processes. While our results point to a lower cost of activity, they do not allow us to address which aspects of network activity decrease energetic needs. Synaptic transmission incurs a greater metabolic cost than action potentials in active networks^34,35^, positioning them as a critical site for reducing energy consumption. Indeed, NMDA-glutamate receptors, which are involved in generating respiratory motor bursts, maintain normal postsynaptic current amplitude after hibernation but adopt a receptor profile that is less permeable to calcium and strongly desensitize, suggesting a lower cost of ion regulation^23^.

We also observed an apparent reduction in the O_2_ consumed by activity-independent processes beyond those contributing to spontaneous synaptic transmission, the Na^+^-K^+^ ATPase, and the V-ATPase. As we could not comprehensively block all additional activity-independent housekeeping process within the brain, applying cyanide to stop remaining respiration allowed us to assess potential reduction in energy use that was not captured with inhibitors of specific physiological processes. Therefore, a smaller change in tissue O_2_ upon the application of cyanide suggests less O_2_ consumption for activity-independent processes in hibernators. There are two non-mutually exclusive possibilities that may explain this finding. First, hibernation may reduce the energy demands of housekeeping processes such as lipid turnover, cellular cargo transport, and other maintenance functions, each of which can incur substantial energy costs^24,36^. Second, these results may reflect alterations in mitochondrial function per se, such as tighter coupling between electron transport and ATP synthesis due to reduced proton leak across the inner membrane, potentially driven by altered uncoupling protein function, membrane lipid composition, or cristae structure^37–41^. Therefore, future work directly assessing mitochondrial function and energy spent on housekeeping physiology will help to discern between these possibilities. Overall, hibernation leads to a reduction in O_2_ consumed in the brainstem through both activity-dependent and activity-independent physiological processes.

We present three additional lines of evidence that neuronal activity incurs a lower aerobic cost while maintaining normal function. In controls, activity declined with tissue pO_2_ and had a critical threshold where rhythmic output became disrupted. In contrast, hibernators were stable from baseline to anoxia and lacked a measurable p_crit_. This shows that network function persists as aerobic metabolism decreases, and even ceases, corroborating our previous reports^12,13^. Further supporting the reduced reliance on aerobic respiration, increasing activity within the physiological range normally elevates O_2_ consumption, but after hibernation, comparable increases in activity are met without consuming further O_2_. While the specific mechanisms for this are not yet known, we hypothesize that glycolysis may be upregulated to provide ATP that supports the increased demands of activity without supplying end-products to the mitochondria and/or network efficiency increases to such a degree where enhanced activity requires minimal additional ATP input. Finally, we demonstrate that disinhibition leads to substantial oxygen consumption in both controls and hibernators, indicating that although the normal dynamic range of activity is supported with less O_2_ consumption, they retain a capacity to increase aerobic metabolism during pathological hyperexcitability. Yet, following this initial consumption event, hibernators stabilized at higher pO_2_ values than controls while maintaining normal activity. While reductions in GABA’s contribution to rhythm generation likely play a role^14^, these results suggest that activity may be preserved because of its reduced energetic requirements, improving the capacity to “share” energetic resources with the elevated metabolic demands of seizure-like bursts. We present multiple lines of evidence that these circuits shift to consume less O₂, and we suggest that similar adaptations may extend to other species. For example, turtles and diving mammals and birds exhibit neural networks capable of maintaining function under low-oxygen conditions^42,43^, suggesting that energy-efficient network function, as described here, may also apply to species traditionally viewed as hypoxia-tolerant. Overall, these results challenge the assumption that neuronal output is constrained by aerobic respiration, highlighting high-efficiency modes of neural communication as a plastic trait.

One additional intriguing feature of these results is that while activity consumed less O_2_ across a range of intensities (Figure 1-3), hibernator mitochondria do not appear to be dysfunctional since they retain a capacity to aggressively consume O_2_ during pathological hyperexcitability (Figure 4). This indicates the relationship between activity and O_2_ operates with a reduced sensitivity, rather than being eliminated. The sensitivity of mitochondrial respiration to neural activity occurs through calcium-dependent mechanisms^44^. During presynaptic firing, calcium influx from extracellular sources enter mitochondria through mitochondrial calcium uniporter (MCU) to stimulate oxidative phosphorylation. MCU’s high sensitivity for calcium uptake in neurons is controlled by a regulatory protein, mitochondrial calcium uptake 3 (MICU3), where its deletion reduces mitochondrial Ca^2+^ uptake and aerobic metabolism^6,45^. In contrast, MICU1 keeps MCU in a closed state, while MICU2 permits less calcium entry than MICU3, leading to less activity-dependent aerobic respiration^6,46,47^. While factors that control the calcium permeability of the MCU in frogs are not known, it presents a biologically plausible explanation for alterations in the relationship between activity and O_2_ consumption as we show: Hibernation may reduce mitochondrial calcium permeability, which prevents neural activity from increasing aerobic metabolism within the physiological range. However, after crossing a calcium threshold during pathological hyperexcitability, O_2_ consumption accelerates. Future work should address the possibility of plasticity in coupling between activity and calcium-dependent activation of mitochondrial respiration.

## MATERIALS AND METHODS

### Experimental model

All procedures were approved by the Animal Care and Use Committee at the University of Missouri (protocol #39264). Adult female American bullfrogs (*Aquarana catesbeiana*, ∼100 g; n = 51) were obtained from Rana Ranch (Twin Falls, ID, USA) and housed in 20-gallon tanks containing dechlorinated, aerated water at room temperature on a 12-h light/dark cycle. Frogs had access to wet and dry platforms, were fed weekly, along with daily water checks and weekly water changes. Control animals were acclimated for ≥1 week before experiments. A subset of frogs was subjected to a simulated hibernation protocol. Food was withheld and water temperature was decreased from room temperature to 4°C over 10 days in a temperature-controlled chamber. Once at 4°C, a perforated screen was placed below the water surface to enforce continuous submergence while allowing gas exchange at the air-water interface, and frogs were maintained for 4 weeks before experimentation. Water pO_2_ was approximately 155 mmHg in both control (∼22°C) and hibernation (4°C) tanks, consistent with air-equilibrated water. Frogs subjected to the cold-submergence treatment are referred to here as “hibernators,” following prior work showing that similar protocols induces metabolic suppression, consistent with the definitions of hibernation that involve reduced whole-animal metabolism beyond that which is driven by reduced temperature alone^48^.

### Brainstem–Spinal Cord Preparation

Brainstem–spinal cord preparations were obtained as described previously^49^. Frogs were anesthetized with isoflurane (∼1 ml/L; v/v) until the toe-pinch reflex was no longer present, then rapidly decapitated with a guillotine. The head was submerged in ice-cold bullfrog artificial cerebrospinal fluid (aCSF; in mM: 104 NaCl, 4 KCl, 1.4 MgCl_2_, 7.5 glucose, 40 NaHCO_3_, 2.5 CaCl_2_, and 1 NaH_2_PO_4_; bubbled with 98.5% O_2_/ 1.5% CO_2_; pH 7.85). The forebrain was removed, and the brainstem–spinal cord preparation (including the midbrain, brainstem, cut caudal to the hypoglossal nerve roots) was isolated with nerve roots intact and transferred to a recording chamber continuously superfused with oxygenated aCSF. Chamber temperature was maintained at 22°C using a Warner Instruments CL-100 bipolar temperature controller with the thermistor placed adjacent to the tissue. Motor output was recorded from the vagus nerve (cranial nerve X) using glass suction electrodes. Vagus nerve recordings provide a reliable index of the overall motor pattern generated by the respiratory network and has been widely used to assess circuit function in this and similar preparations^15,50,51^. Signals were AC-amplified (1000×, Model 1700; A-M Systems, Carlsborg, WA), filtered (10Hz-5kHz), digitized (PowerLab 8/35, ADInstruments, Colorado Springs, CO), rectified, and integrated online (time constant = 100 ms) to visualize rhythmic motor bursts. Vagus nerve output was allowed to stabilize for ≥1h in aCSF bubbled with 98% O_2_/1.5% CO_2_ prior to any experimental manipulation.

### Oxygen Profiling

Tissue oxygen partial pressure (pO_2_) was measured using a Clark-type microsensor (OX-50; Unisense A/S, Denmark) with a 50 μm tip diameter, defining a local detection volume on the order of tens of microns. The sensor has a response time of ∼5 s, resulting in a ∼5s temporal averaging of the pO_2_ signal with a negligible oxygen consumption relative to tissue, drawing 4×10^-4^-5×10^-3^ nmol O_2_/hr during operation. The sensor was calibrated at 22 °C in air-saturated water and in an anoxic solution (0.1 M sodium ascorbate in 0.1 M NaOH). The detection limit of the OX-50 sensor is ∼0.3 µmol/L (∼0.165 mmHg under our experimental conditions; 22°C, ∼755 mmHg atmospheric pressure), with a noise floor of approximately 0.2 mmHg. The analog signal from a PA2000 amplifier (Unisense A/S) was digitized through an ADC-216USB converter. For all experiments, the sensor was inserted slightly lateral to the midline of the brainstem in between the vagus and trigeminal nerves. The tissue pO_2_ was recorded in 50 μm vertical steps from the surface to a depth of 350 μm, an area that encompasses part of the respiratory network^52^. At each 50 μm vertical step, the sensor was allowed to stabilize for 20 seconds before recording the pO_2_ value, which was taken as the tissue pO_2_ at that depth. In 6 experiments for each group, the sensor was advanced beyond 350 µM to up to 650 μm, near the center of the tissue.

The amphibian respiratory network tends to produce rhythmic output with bursts often clustered into stronger multi-breath episodes but can also assume a variety of output patterns^51^. While the respiratory network is distributed throughout the brainstem^52^, tissue pO_2_ fluctuations detected by the microsensor typically occurred in phase with “crescendo-like” breathing episodes when these occurred (Figure 1A), confirming that the sensor reported oxygen dynamics within or around the region of the active respiratory network. The sensor was only able to detect oxygen fluctuations associated with these high-drive episodes that spanned the 5 s detection duration of the sensor, as individual, single bursts that are most dominant in this preparation are 1s in duration, and therefore, likely produce oxygen transients that are faster than the response time of the sensor. Because the oxygen microsensor reports pO_2_ from spatially limited region around the tip, we also cannot assign the precise anatomical compartment neurons contributing to each measurement (*e.g*., in regions densely packed with cell bodies, synapses, or a combination of the two). The electrode tip may lie closer to somatic regions, axon fiber tracts, or synaptically dense neuropil, all of which have the potential for different degrees of oxygen consumption^53^. Thus, the recorded pO_2_ reflects the integrated activity of multiple cellular processes within that microenvironment, which was recorded as consistently as possible in the same location across experiments. Targeted assessments of oxygen consumption in sub-cellular compartments require further investigation.

### Pharmacological Experiments

To estimate the contribution of aerobic metabolism to different ATP-consuming processes, inhibitors specific to each process were sequentially blocked while monitoring tissue pO at a fixed electrode depth (350 μm). Drugs were applied cumulatively, i.e., each subsequent inhibitor was added to the superfusate without washing out the previous drug(s). The following final bath concentrations were applied in sequence: Tetrodotoxin (TTX, voltage-gated Na⁺ channel blocker, 500nM); DNQX+APV (AMPA receptor antagonist, 10μM; APV (NMDA receptor antagonist, 50μM); Bafilomycin A1 (V-ATPase inhibitor, 1μM); Ouabain (Na⁺/K⁺-ATPase pump inhibitor, 500μM); Sodium cyanide (mitochondrial Complex IV inhibitor, 5mM). Each drug was superfused until a stable plateau in pO_2_ was achieved (∼15–20 min), and then the next was applied. Because all previously applied drugs remained in the bath, the ΔpO_2_ following each new inhibitor (the steady-state pO_2_ reading before the drug subtracted from the steady-state reading after) reflects the loss of the O_2_ that was being consumed specifically by the newly blocked process, serving as a functional readout of the relative metabolic contribution of the targeted pathway. Seven control and seven hibernated brainstems were used. Although our pharmacological approach isolates specific pathways powered by aerobic metabolism, the relative magnitude of their pO_2_ changes depends on their local abundance within the sampled volume. As a result, oxygen consumption associated with spontaneous synaptic transmission or vesicle recycling may be underestimated if the sensor was positioned nearer to neuronal cell bodies than to synapse-rich zones^53^. These factors are inherent to single-point oxygen microsensors and reflect the spatial averaging of pO_2_ over the sensor’s diffusion field and require further assessments targeting specific sub-cellular recordings.

### Graded Hypoxia and Increased Activity

Graded hypoxia was induced by progressively replacing O_2_ with N_2_ in the aCSF using a mass-flow controller (MFC-4, Sable Systems International). The N_2_ fraction was increased by 10% every 10 min until anoxia (0% O_2_ in the reservoir) was reached. When bubbled with 98.5% N_2_/ 1.5% CO_2_, bath pO_2_ surrounding the tissue was ∼15 mmHg due to mixing at the liquid-air interface. In all controls and hibernators, tissue pO_2_ at the sensor reported 0 mmHg; thus, we refer this outcome as “anoxia.” Therefore, while the sensor reported 0 mmHg inside the tissue when bathed with solution gased with 98.5% N_2_/ 1.5% CO_2_ (and in some cases at even higher fractions of O_2_), it is not possible to resolve differences in tissue pO_2_ below the sensor’s detection limit (see Oxygen Profiling). Regardless, a zero reading under these conditions reflects an infinitesimal amount of O_2_ should any exist in the tissue. Six control and six hibernated brainstems were tested in this series. Tissue pO_2_ and respiratory burst frequency were recorded continuously (n = 6 controls; n = 6 hibernators). We interpret p_crit_ as the O_2_ tension where oxidative phosphorylation decreases to a point that can no longer preserve normal neural activity, beyond which activity was either chaotic or irregular.

To assess the metabolic cost of elevated physiological activity, isoproterenol (10 μM) was bath-applied after ≥1 h of stable baseline activity and superfused for 30min (n=7 controls; n=6 hibernators). While other stimuli such as hypercapnia could be used to stimulate respiratory frequency, in our hands, the responses can be inconsistent from preparation to preparation, while isoproterenol produces consistent increases in frequency. To evaluate responses to pathological hyperexcitability, bicuculline methiodide (10 μM) was superfused for 30 min to induce non-respiratory, seizure-like bursts (n=6 per group). Non-respiratory bursts were manually counted based on their prolonged duration and distinct waveform morphology, as previously described^28^. The oxygen microsensor integrates over ∼5s which does not resolve the rapid pO_2_ transients associated with individual respiratory bursts (∼1s duration). Therefore, pO_2_ data for these experiments reflect longer-timescale changes in tissue oxygen dynamics, rather than instantaneous fluctuations tied to single bursts. This averaging is appropriate for quantifying sustained differences in oxygen consumption over minutes, which were apparent in these experiments.

### Drugs

Tetrodotoxin (TTX), DNQX, APV, and bicuculline were obtained from Hello Bio. Bafilomycin A1 was purchased from Cayman Chemical. Ouabain was obtained from Thermo Scientific, sodium cyanide from MP Biomedicals, and isoproterenol from Tokyo Chemical Industry. All drugs were prepared as stock solutions according to manufacturer recommendations and diluted to final concentrations in artificial cerebrospinal fluid immediately prior to use.

### Data Analysis

Respiratory burst frequency was determined from rectified and integrated vagus recordings using the Peak Analysis function in LabChart 8 (ADInstruments). Bursts were identified by standard criteria (∼1 s duration) with start/stop thresholds at 5% of peak height^50^. For graded hypoxia experiments, mean burst frequency and tissue pO_2_ were calculated for every 10 mmHg drop in tissue pO_2_ relative to baseline at 98% O_2_. For experiments involving sequential drug exposures, mean pO_2_ values were obtained over the final 10 min of each treatment at steady-state. During the application of isoproterenol, pO_2_ values were taken at baseline and then at 10 minute intervals sampling for 2 minutes during the 30 minute exposure. For bicuculine, pO_2_ values were taken at baseline, at the trough of the large dip caused by the large non-respiratory burst following bicuculine, and then the last 5 minutes of the 30-minute exposure

### Statistical analysis

Data are presented as mean ± SD or box and whisker plots unless otherwise stated. Statistical comparisons were performed in GraphPad Prism v10.6.1 (San Diego, CA, USA). Two-way repeated measures ANOVA was applied with two independent variables and one dependent variable (e.g., the effect of depth and group on tissue pO_2_). In these cases, Holm–Šidák multiple comparisons test was applied after if there were any interaction or group effects. To assess the effects of tissue pO_2_ and burst frequency, we used a mixed linear model with restricted maximum likelihood, as this dataset had many missing values due to the differing starting points of pO_2_ from preparation to preparation. Unpaired t tests were used to compare two independent groups. Individual data points are shown where appropriate to show variability among preparations.

## CONCLUSION

Our findings reveal that neural circuits in the vertebrate brain can undergo a fundamental reorganization of the relationship between activity and O_2_ consumption. By reducing the aerobic requirement for neural activity, the frog brainstem maintains stable network output under metabolic conditions that normally induce severe perturbations. The relevance of this reduced aerobic cost is likely for emergence from hibernation, when frogs must restart the neural circuits that drive breathing before the first breath is taken, at which point brain oxygenation has not yet been restored by lung ventilation. A central element of this shift appears to be the enhanced efficiency of network function, which allows hibernators to generate normal output with less ongoing aerobic metabolism. These results raise the question of how neural circuits can transition between vastly different aerobic requirements while maintaining similar functional output. Understanding these mechanisms may be particularly relevant for neurological disorders such as Alzheimer’s and Parkinson’s disease, where impaired aerobic metabolism compromises circuit function^8^. By identifying mechanisms that allow circuits transition between states with vastly different aerobic demands needed to produce similar function, our work highlights how natural adaptation in diverse animal models can serve as valuable frameworks for exploring strategies to improve metabolic resilience in the mammalian brain.

## Acknowledgements

The authors would like to thank Nik Bueschke and Jose Viteri for comments on the manuscript.

