## Supplemental Information for "Frogs uncouple neural activity from oxygen consumption after hibernation"

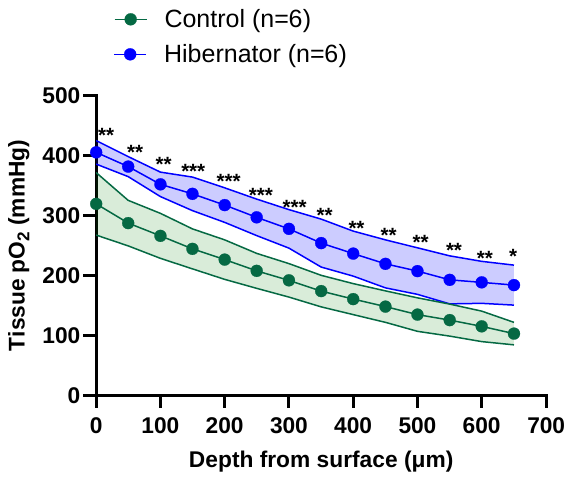

**Supplemental Figure 1 pO_2_ depth profile to 650 μm below the brainstem surface.** Mixed effects model. Main effect of group: p=0.0006, main effect of depth: p<0.0001; interaction, p=0.3866. Holm-Šídák's multiple comparisons test, ***p<0.001, **p<0.01, *<0.05. Group effect manifests as higher pO_2_ in hibernators compared to controls at greater distances from the surface. However, the overall shape of the depth vs. pO_2_ response was similar; thus, no interaction was observed.

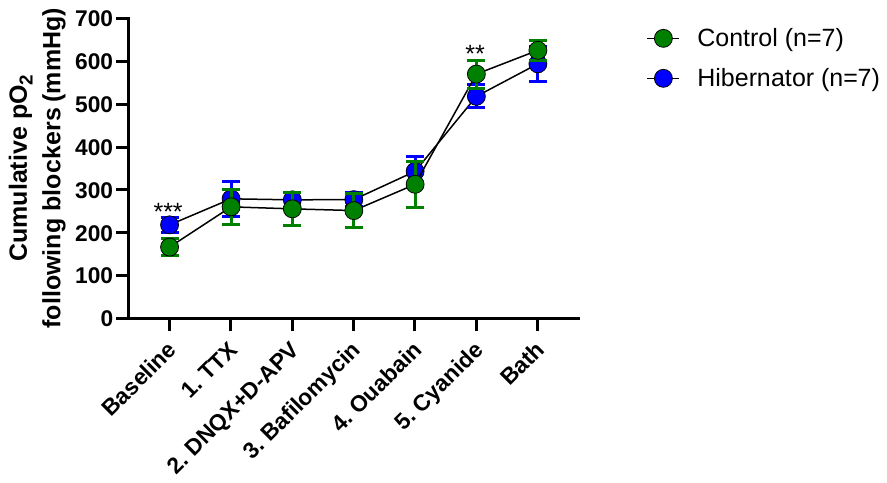

**Supplemental Figure 2 Absolute changes in tissue pO_2_ during sequential drug applications to block different physiological processes.** Raw pO_2_ data used to calculate the change in pO_2_ in response to each drug shown in Figure 1F. Repeated measures two-way ANOVA. Main effect of drug applications: p<0.0001; main effect of group: p=0.5271; interaction between group and drug applications: p<0.0001. Holm-Šídák's multiple comparisons test ***p<0.001, **p<0.01). Interaction arises because hibernators start from a significantly higher baseline pO_2_ and a smaller change in response to cyanide application. These results are consistent with those expressed as a change in pO_2_ following each drug shown in Figure 1F.
